# Metadata Collector: An Open-Source Platform for Standardized Metadata Management in Multi Centre Sequencing Projects

**DOI:** 10.64898/2026.06.05.730314

**Authors:** Renato Liguori, Michael Huttner, Fulvia Ferrazzi

**Affiliations:** Department of Nephropathology, Institute of Pathology, Friedrich-Alexander-Universität Erlangen-Nürnberg, Krankenhausstrase 8-10, Erlangen, 91054, Bavaria, Germany; Institute of Pathology, Friedrich-Alexander-Universität Erlangen-Nürnberg, Krankenhausstrase 8-10, Erlangen, 91054, Bavaria, Germany; Faculty of Informatics and Data Science (FIDS) University of Regensburg, Am Biopark 9, Regensburg, 93053, Bavaria, Germany

## Abstract

**Background:** Next-generation sequencing (NGS) projects generate increasingly complex metadata that are critical for reproducibility, interoperability, and compliance with FAIR principles. Nevertheless, metadata curation in multi-institutional settings often still relies on spreadsheets, manual data entry and curation, as well as non-standardized terminology. These practices frequently result in incomplete or inconsistent annotations, hinder metadata sharing, and delay submission to public repositories.

**Results:** We developed Metadata Collector as a React/API/PostgreSQL web platform and deployed it on a Kubernetes cluster within a large German research consortium. The platform implements a flexible, machine-readable metadata model for experimental data and integrates customizable templates, controlled vocabularies designed to support future ontology integration, and a complete event-based versioning model. Since deployment, Metadata Collector has been used across 32 projects involving RNA-seq, scRNA-seq, ATAC-seq and multiomics datasets, representing over 700 annotated samples contributed by multiple consortium partners. The platform is designed for use by non-computational researchers as well as centralized facilities and can be integrated into existing research data management infrastructures.

**Conclusions:** Metadata Collector embeds standardization early in the metadata lifecycle, ensuring consistent, FAIR-aligned, and reproducible metadata across distributed research groups. Its modular, open-source architecture supports both local and consortium-scale deployments and provides a foundation for future extensions, including multi-omics support and integration with laboratory information management systems and automated submission pipelines.

**Availability:** Open-source under MIT license: https://github.com/spang-lab/metadata-collector

Graphical abstract of Metadata Collector

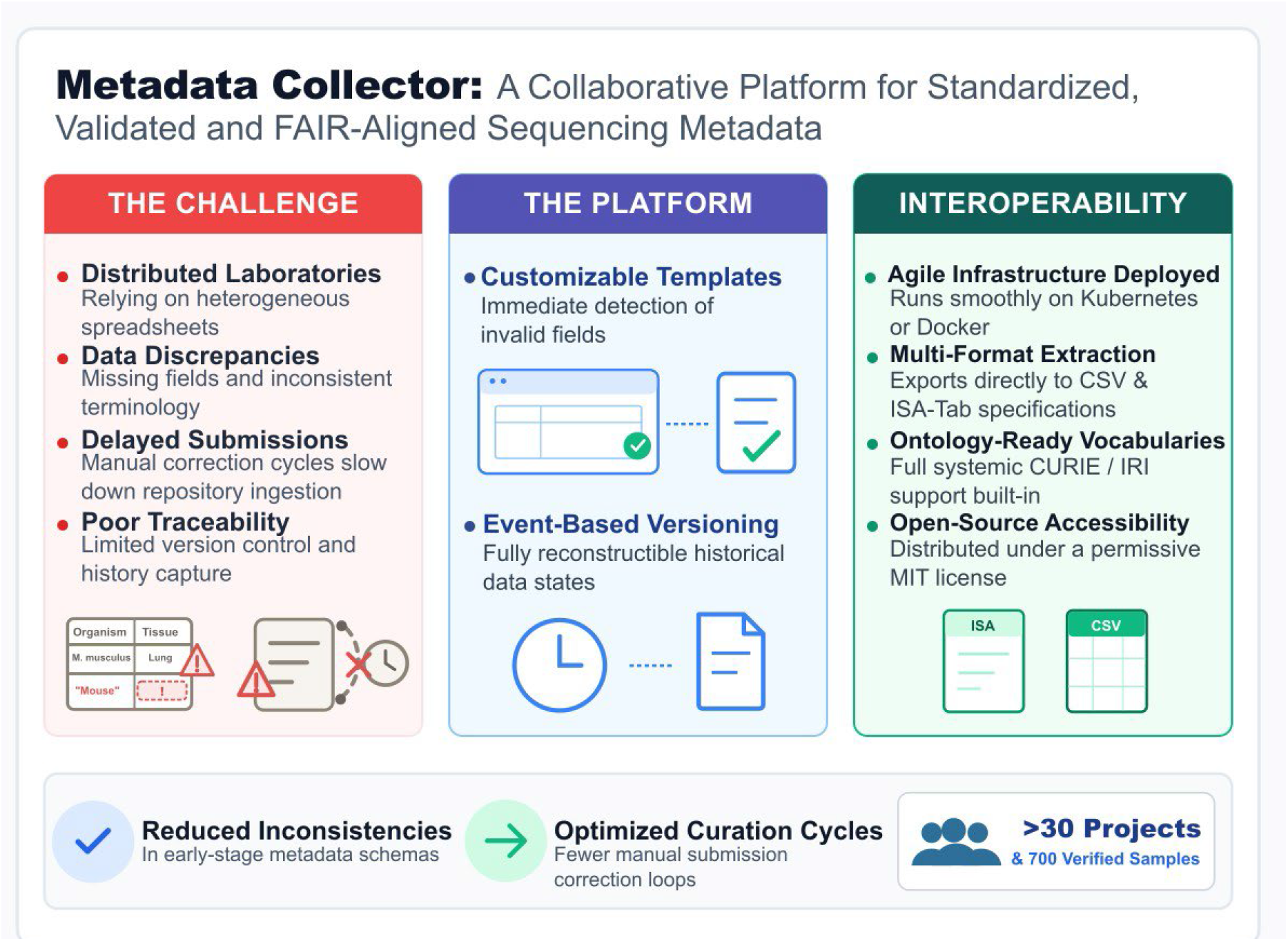

## Introduction

High-quality metadata are essential for ensuring the reproducibility, interoperability, and long-term usability of next-generation sequencing (NGS) data (1, 2). As multi-centre biomedical consortia increasingly generate large, heterogeneous sequencing datasets, the challenge of collecting accurate, consistent, and FAIR-aligned metadata has become more pronounced (3, 4). Metadata originate from diverse laboratories, each with its own conventions, spreadsheets, and documentation practices, often resulting in fragmented annotation, inconsistent terminology, and need for extensive manual curation. These issues affect the efficient use of analysis pipelines, repository submissions, and ultimately the reusability of the generated datasets.

A wide range of tools and standards has emerged to support different stages of the metadata lifecycle, yet they only partially address the challenges faced by multi-centre consortia. The ISA framework (ISA-Tab and ISA-JSON) provides a flexible representation for experimental metadata and promotes the use of controlled vocabularies (5). However, its primary focus is on metadata representation and exchange, offering limited support for collaborative metadata management across distributed research groups. Alternative ecosystems like the CEDAR Workbench have attempted to bridge this template authoring gap (6); however repository submission systems, such as ENA, NCBI BioSample, and EGA, still enforce metadata requirements by validating mandatory fields and formatting during data submission (7-10). While essential for repository compliance, these systems operate primarily at the final stage of the data lifecycle and provide limited support for the collaborative, iterative, and often error-prone metadata creation process that occurs upstream during sample collection and project coordination.

Laboratory Information Management Systems (LIMS) and electronic laboratory notebooks (ELNs) (11) offer deeper integration with laboratory workflows and sample tracking. However, their deployment often requires substantial customization, dedicated infrastructure, and institutional support, which can limit their adoption as shared metadata management platforms across independent research groups. As a result, many consortia continue to rely on spreadsheets and ad hoc solutions for metadata collection and harmonization because of their accessibility and flexibility (12). While convenient, spreadsheet-based approaches lack critical features for multi-centre coordination, including version control, access management, controlled vocabularies, real-time validation, and transparent tracking of changes. Consequently, metadata management often becomes fragmented, leading to duplicated effort, ambiguous terminology, conflicting record versions, and repeated rounds of manual correction before data can be analysed or submitted to public repositories.

Efforts to improve metadata interoperability, such as OBO Foundry ontologies (13), EDAM (14), and domain-specific curated vocabularies, provide valuable resources for standardizing terminology. However, these standards are rarely integrated into user-friendly platforms that actively support metadata entry and harmonization. As a result, even when standardized vocabularies are available, their adoption remains inconsistent unless users are guided during the annotation process. This disconnect frequently contributes to downstream bottlenecks, including metadata inconsistencies, multiple submission revision cycles, and delays in dataset release.

To address these challenges, we developed Metadata Collector, a web-based platform designed to support standardized, validated, and collaborative metadata management throughout the lifecycle of NGS projects. The platform aims to reduce annotation inconsistencies at their source, streamline communication among distributed research groups, and ensure that metadata remain consistent, traceable, and aligned with FAIR principles from project initiation to public data submission.

## Material and Methods

### System Architecture Overview

Metadata Collector follows a modular client–server architecture designed to support efficient interaction, scalable deployment, and maintainable long-term operation in multi-centre environments. The frontend is implemented as a single-page application (SPA) that communicates with the backend through RESTful API endpoints (Figure 1). This separation ensures a clear boundary between presentation, business logic, and data management, enabling flexible deployment across institutional infrastructures.

**Figure 1:**
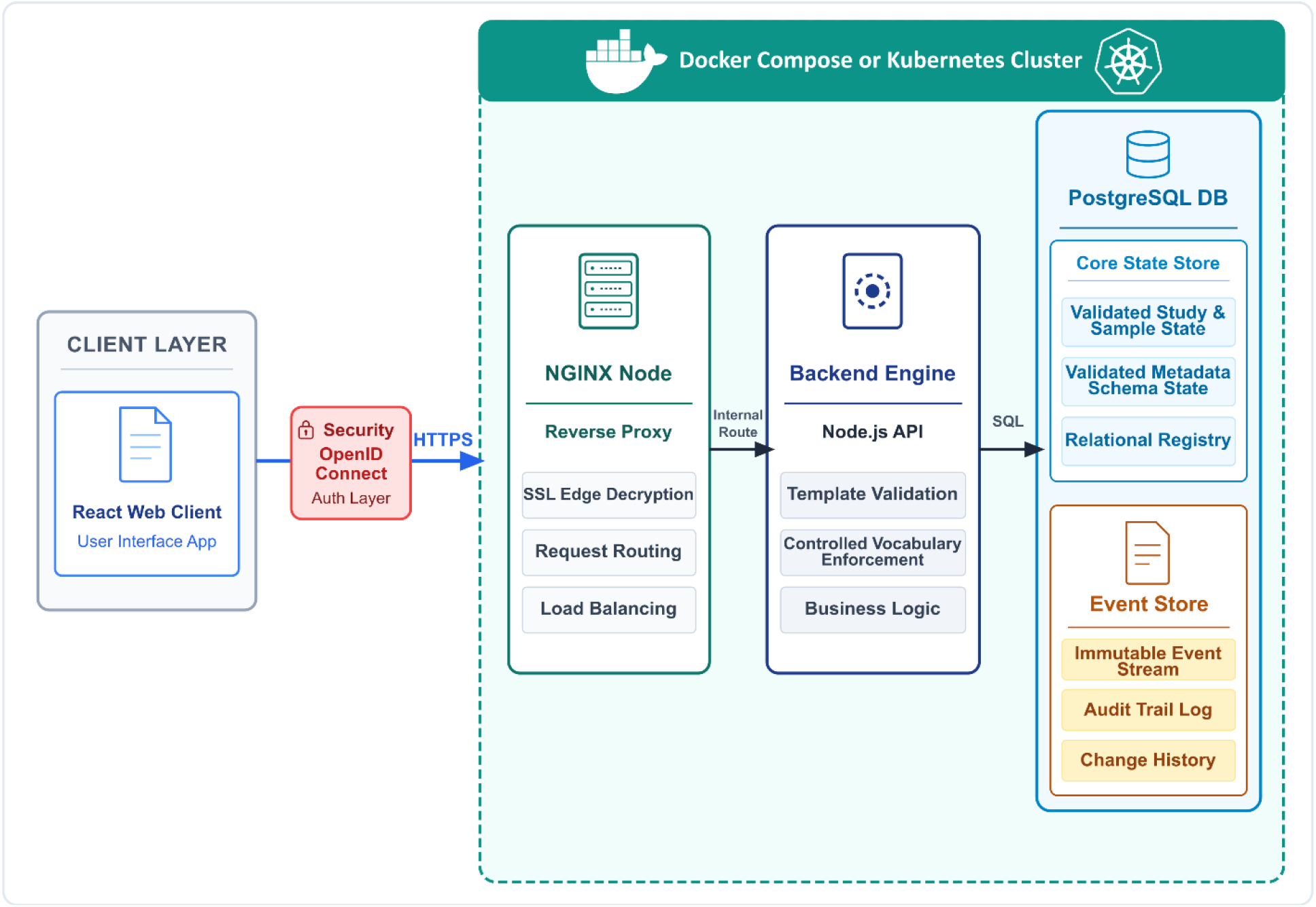
System architecture and containerized cluster deployment of Metadata Collector. Users interact with a React-based web interface connected via HTTPS/TLS to a containerized deployment environment. Incoming requests are routed through an NGINX reverse proxy to the backend validation and business logic layer, where template validation and controlled vocabulary enforcement are performed. Validated metadata are stored in a PostgreSQL state database, while all changes are recorded in a separate append-only event store to support versioning and auditability.

The backend exposes endpoints for project creation, metadata editing, validation, controlled vocabulary management, and event-based versioning. All modifications are stored as immutable events in an append-only log, from which the current metadata state is reconstructed. This event sourcing architectural pattern avoids the pitfalls of schema inflation or ad hoc audit tables while guaranteeing complete auditability (15). Metadata and events are stored in PostgreSQL, with separate tables for the event store and derived state to support fast queries and full historical recovery.

### Technology Stack and Deployment Environment

The reference deployment of Metadata Collector uses a React-based client application and a Nodejs backend API (Figure 1). PostgreSQL serves as the metadata and event-state database. All components are containerized and orchestrated through Kubernetes, enabling load distribution, horizontal scaling, rolling updates, and centralized monitoring (e.g., via Prometheus and Grafana). Static assets are served through a NGINX reverse proxy that handles routing, compression, and TLS termination.

API responses are optimized through asynchronous endpoints and connection pooling, resulting in low latency (typically <50 ms for common operations in consortium-scale deployments). For smaller projects or local development, Metadata Collector can be deployed via a Docker Compose configuration that includes all core services with minimal external dependencies. This setup mirrors the production container structure, ensuring consistent behaviour between local and cluster-based deployments (Figure 1).

Exact software versions are tracked and documented in the public GitHub repository to ensure reproducibility while allowing continuous updates.

### Security and Data Protection

Security and data protection were central design considerations due to the multi-institutional nature of the platform. Authentication is handled through OpenID Connect, allowing integration with institutional identity providers. User roles and permissions are enforced at the platform level, ensuring that only authorized contributors can view, edit, or export specific metadata. All communication is secured via TLS encryption. Within the Kubernetes cluster, network policies restrict pod-to-pod communication, ensuring that the PostgreSQL database and event store are only accessible from backend services. No direct database access is exposed to users.

Metadata contain no direct personal identifiers: all entries are pseudonymized and linked to study-level identifiers rather than individuals, facilitating compliance with institutional data protection policies and GDPR requirements. Passwords, tokens, and session information are not stored in the database.

### Data Model

The data model is structured around three abstractions: entities, properties, and events, all managed in PostgreSQL.

- *Entities* represent the main metadata objects. Currently, two entity types are implemented: *Projects* and *Samples*, with *Samples* always hierarchically linked to their parent *Project*.
- *Properties* correspond to metadata fields, analogous to spreadsheet columns, which can be associated with entities. This flexible design allows both mandatory and optional fields to be defined and extended according to evolving assay requirements.
- *Events* capture every user interaction with the system. Instead of overwriting existing values, edits are stored as discrete events in a chronological log. This ensures that no information is lost and provides a complete version history, allowing reconstruction of any previous project state at a chosen time point; on our reference deployment, state reconstruction for consortium-scale datasets (≤700 samples) typically completes within milliseconds on the reference deployment.

Controlled vocabularies are stored in dedicated tables and can include machine-readable identifiers (e.g., CURIEs or IRIs), enabling future linkage to external ontology services. By combining entity-based organization, property-driven flexibility, and event-based traceability, the data model supports both robust governance (immutability, auditability) and adaptability to new metadata requirements.

### Client Application

The user interface is implemented as a single-page web application (SPA) developed with the React framework. The client interacts exclusively with the server API, dynamically rendering metadata tables and forms. Each user action, such as adding a sample, editing a field, or exporting metadata, is transmitted as a discrete event to the backend, ensuring complete traceability. By separating data handling from the interface, the architecture supports modular development and facilitates future integration with third-party applications.

### Home Interface and Project Overview

Upon login to Metadata Collector, users are presented with the home interface, which serves as the central hub for all metadata management activities. The dashboard lists all active projects, showing key information such as the project’s name, owner’s contact details (e.g., email) and creation date, allowing users to navigate efficiently between studies.

### Metadata Templates

Templates define the structure and constraints of metadata within each project and serve as schema artefacts specifying field types, as well required vs optional fields. Two main template types are supported:

- predefined consortium templates containing standardized metadata fields across research groups to capture both experimental context and key sample characteristics, aligned with repository and consortium requirements.
- custom templates configured to match specific needs or unique requirements of individual studies.

Templates are stored as JSON descriptors in the database and loaded dynamically by the client application, ensuring that all contributors operate under a shared, consistent schema.

Users with administrative privileges can create or modify templates through a dedicated configuration panel. This GUI translates template definitions into structured forms, allowing administrators to add fields, assign data types, select vocabularies, or adjust validation logic without directly editing configuration files. This approach guarantees schema integrity while allowing templates to evolve during project development.

### Metadata Entry and Validation

During metadata entry, the client application constructs dynamic forms from the active template, applying validation logic (e.g., allowed values, required fields) in real time. The frontend performs preliminary client-side checks, while the backend enforces definitive server-side validation before events are accepted. This dual-layer strategy improves responsiveness while preventing invalid or incomplete submissions.

Each project acts as a structured container for high-level study metadata, providing context for all associated samples and sequencing data. A core set of project-level fields is mandatory (e.g., project name, brief description, principal contributors, overall study design), ensuring clarity and reproducibility. Samples are then added under the project using forms that distinguish between:

- mandatory fields capturing essential identifiers and descriptors required for downstream analysis and repository submission (e.g., Sample ID, Organism, Tissue Type, Disease Group, Cell Population, Sample Type).
- optional fields adding scientific and experimental context (e.g., Subject ID, Phenotype, Treatment Protocol, Library Strategy, and assay-specific processing parameters for, e.g., RNA-seq, ATAC-seq, or spatial transcriptomics).

To promote consistency, many fields are populated via dropdown menus backed by controlled vocabularies, while free-text fields are available for rare or study-specific values. This combination of structured input and flexibility balances standardization with adaptability.

Completed metadata can be exported in multiple formats, such as CSV and ISA-Tab, to ensure compatibility with public repositories (e.g., EGA, SRA) and downstream analysis workflows. An access-controlled audit panel supports collaborative metadata editing across institutions while preserving full traceability of changes.

### Deployment and Infrastructure

To support both production-scale multi-centre deployments and lightweight testing or development setups, Metadata Collector is distributed in two deployment modes both relying on containerized components for reproducibility and ease of maintenance, a Kubernetes Deployment and a Docker Compose Deployment.

#### Kubernetes Deployment

The reference production deployment uses Kubernetes to orchestrate the application stack, including the frontend, backend, and PostgreSQL database. Helm charts and deployment manifests define the configuration of each service, enabling automated scaling, rolling updates, health checks, and resource allocation. The platform integrates with institutional identity providers through OpenID Connect and leverages Kubernetes networking policies to restrict internal traffic to authorized services making this mode suitable for institutional clusters and consortia requiring high availability and secure multi-tenant operation.

#### Docker Compose Deployment

For smaller projects, testing environments, or local development, Metadata Collector can be deployed via a single Docker Compose configuration. This setup includes all core services, React frontend, Node.js backend, and PostgreSQL database, preconfigured with minimal requirements and no external dependencies. By mirroring the production container structure, the Docker Compose mode ensures consistent behaviour between local and cluster deployments, simplifying development and onboarding.

### Open-Source Distribution

The full codebase, including deployment manifests, is openly available under an MIT license on GitHub. Container images and versioned releases are published on public registries to facilitate reproducible installations.

## Results

### Key Features of Metadata Collector

Metadata Collector provides a set of core functionalities that support consistent, validated, and collaborative metadata management in multi-centre sequencing projects.

#### Key capabilities include

- Customizable metadata templates Project- or assay-specific schemas define field types, controlled vocabularies, and validation rules, ensuring structural consistency while accommodating diverse experimental designs.
- Real time validation Client-side checks and server-side enforcement prevent incomplete or invalid entries, reducing downstream curation effort.
- Controlled vocabularies Dropdown lists and ontology-ready term management promote standardized terminology across research groups.
- Event-based versioning and traceability All edits are stored as discrete events, allowing complete reconstruction of prior states and transparent tracking of contributions.
- Collaborative editing with role-based access control Multi-user, permission-aware workflows enable distributed teams to contribute metadata while preserving auditability and ownership.
- Interoperable export formats Metadata can be exported in CSV and ISA-Tab formats, supporting submission to public repositories (e.g., EGA, ENA) and integration with analysis pipelines.

Together, these features provide the foundation for the platform’s adoption and its alignment with the FAIR principles, as described in the following sections.

### Workflow and FAIR Principles Alignment

The platform incorporates several features intended to support FAIR-aligned metadata management. To support findability, metadata are organised through structured framework supported by integrated search functionality. Each project and sample are assigned a unique, persistent internal identifier, enabling unambiguous referencing throughout the platform (Figure 2). Versioning preserves that all historical states remain discoverable, while role-based permissions guarantee that users see only records relevant to their projects (Figure 2). Metadata accessibility is ensured through authenticated web interfaces and standardized export options. Access control is implemented via OpenID Connect and role-based permissions, enabling secure cross-institutional collaboration without exposing sensitive information. Metadata can be exported in open, widely used formats such as CSV and ISA-Tab, facilitating retrieval and downstream use (Figure 2). Because metadata contain no direct personal identifiers and are pseudonymized at source, authorized collaborators can access records without introducing privacy risks.

**Figure 2:**
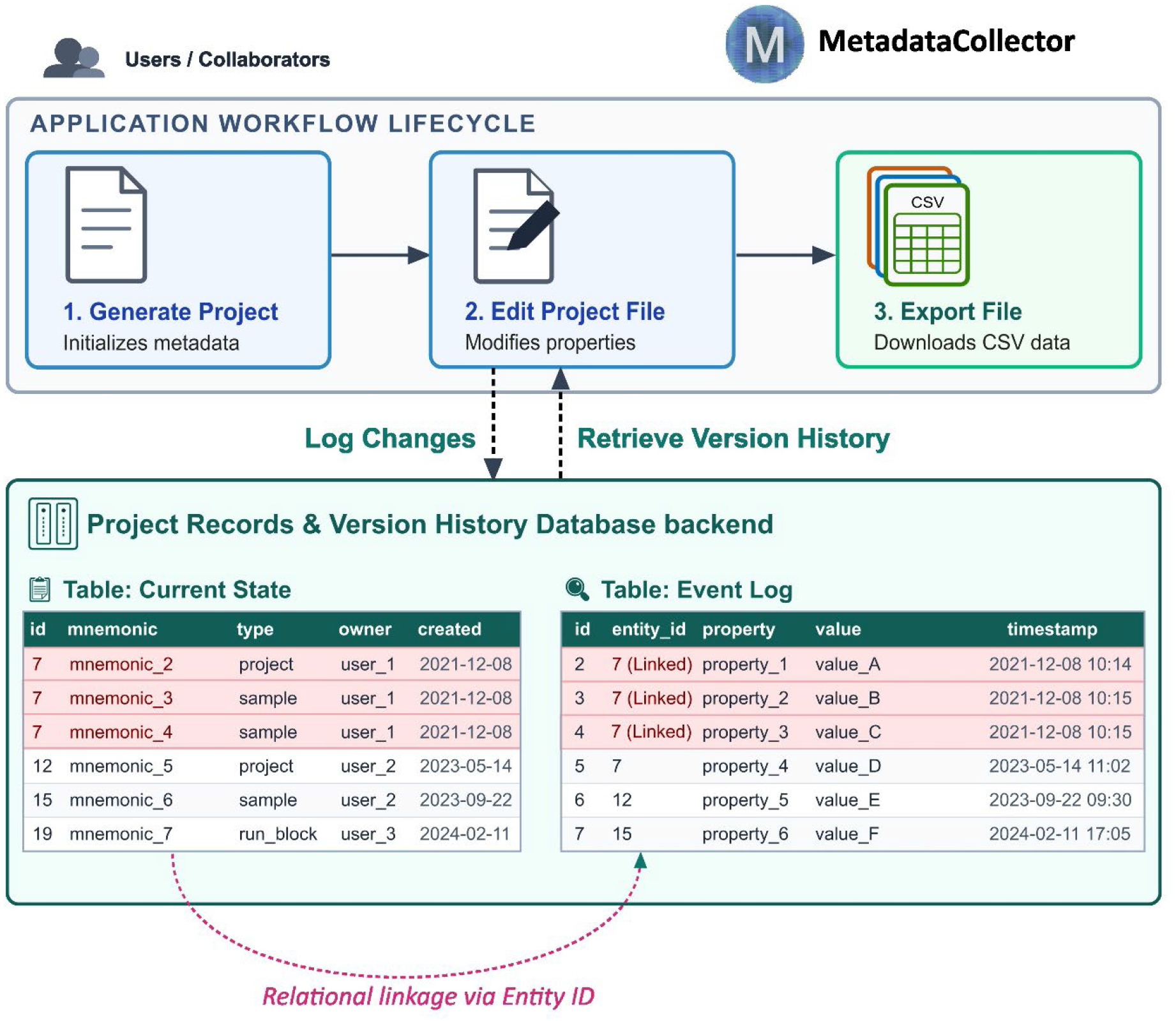
Metadata Collector workflow and event-driven data model. Users create, edit, and export metadata records through a structured workflow. The backend maintains the current metadata state in a project records table while simultaneously recording all modifications in an append only change log linked through entity identifiers. This separation supports metadata provenance, versioning, auditability, and reconstruction of historical record state.

Interoperability is facilitated through controlled vocabularies, structured templates aligned with repository requirements, and export formats compatible with ENA, EGA, and related resources. Reusability is enhanced through rich contextual metadata, transparent provenance, and complete version histories (Figure 2). Mandatory project-level fields capture study design, contributors, and experimental objectives, while sample-level fields record the biological and technical descriptors needed for downstream analysis or future reuse. All modifications are stored as a time-stamped, user-attributed record, providing a complete and traceable history of the metadata. Standardized validation rules further minimize inconsistencies that could hinder downstream integration, thereby enhancing the reliability and reusability of curated metadata across analytical pipelines and data repositories.

### Platform Adoption and Practical Impact

Since its deployment within a large research Consortium, namely the TRR 305 consortium “*Striking a moving target: From mechanisms of metastatic colonization to novel systemic therapies*” funded by the German Research Foundation, the Metadata Collector has been adopted in more than 30 projects, supporting more than 700 samples spanning from RNA-seq, scRNA-seq, ATAC-seq, and multiomics assays.

Table 1 summarizes the principal features of Metadata Collector and compares them with commonly used metadata management solutions. In contrast to existing approaches, which typically perform validation and consistency checks only during repository submission or within individual workflow environments, Metadata Collector incorporates template-driven validation, controlled vocabularies, and record-level versioning directly into the metadata creation process. This integrated approach enables collaborative, real time harmonization of metadata across distributed research groups, improving consistency, traceability, and compliance with repository requirements before downstream analysis or data submission. As a result, metadata quality is addressed proactively rather than through retrospective curation, reducing the effort required for subsequent correction and harmonization.

**Table 1.** High-level feature comparison of Metadata Collector with commonly used metadata formats and platforms. The comparison is based on publicly documented functionality and typical usage patterns and is intended to highlight conceptual and functional differences rather than provide an exhaustive technical benchmark. Features described as “External” or “Platform-dependent” reflect typical usage in common deployments and may vary depending on local configurations, extensions, or institutional implementations.

| *Feature* | *Metadata Collector* | *ISA-Tab / ISA Tools* | *Galaxy* | *NCBI BioSample* | *Custom Spreadsheets* |
| --- | --- | --- | --- | --- | --- |
| **Real time validation** | **Supported**<br>(Template- and vocabulary-based) | **On-demand**<br>(Via external validators) | **Execution-time**<br>(Tool/datatype checks) | **Post-upload**<br>(Batch submission-time) | **Local only**<br>(Basic cell formulas) |
| **Customizable templates** | **Supported** (GUI-based) | **Supported**<br>(Configurator-driven) | **Constrained**<br>(Tool-specific wrappers) | **Restricted**<br>(Predefined Attribute Packages) | **Ad-hoc**<br>(Manual layout) |
| **Multi-user metadata editing** | **Granular** (RBAC) | **External**<br>(Repository-dependent) | **Only histories and workspace sharing** | **Not supported** | **File-sharing dependent**<br>(e.g., OneDrive) |
| **Version control** | **Record-level**<br>(Event-driven) | **External**<br>(VCS/Git-dependent) | **Workflow provenance**<br>(Dataset-level)* | **Not supported** | **File-level**<br>(Cloud version history) |
| **Audit trail** | **Granular** (Edit-level logging) | **External** (Git commit logs) | **Workflow-level**<br>(Provenance history)* | **Submission logs** (Receipt only) | **Minimal**<br>(Basic file metadata) |
| **Export formats** | **CSV, ISA-Tab** | **ISA-Tab, ISA-JSON** | **TSV/JSON**<br>(ISA via tools) | **XML, JSON, TSV** | **Not standardized** |
| **Open source** | **Yes** | **Yes** | **Yes** | **No** | **N/A** |

\*Galaxy version control and Audit trail refers to workflow execution level rather than record level metadata versioning and step-by-step edit log. RBAC = Role-Based Access Control

Consortium partners reported a substantial reduction in annotation inconsistencies compared with spreadsheet-based workflows. In several representative projects, initial data imports generated validation warnings, most commonly due to missing mandatory fields. These issues were resolved early in the metadata lifecycle, rather than during later submission or analysis stages. Consequently, fewer manual correction cycles were required prior to repository submission compared with pre-platform workflows. Terminology inconsistencies, such as heterogeneous free-text entries for tissue type or organism, also declined markedly after adoption of controlled vocabularies embedded within shared templates. Consortium members also reported a reduction in internal email exchanges and manual revision loops, as communication and coordination were centralized around a shared, versioned metadata workspace

### Use Case Demonstration: Managing a Consortium-Scale NGS Project

Metadata Collector supports guided metadata creation and harmonization through an interactive web interface designed for distributed research teams with heterogeneous technical backgrounds. To illustrate the platform usage, we describe a representative workflow based on common metadata management scenarios within TRR305.

The workflow begins with project initialization, where users define the study metadata and select an appropriate template that determines the structure of subsequent sample annotation (Figure 3). Researchers then enter study-level metadata through structured forms automatically generated from the active template. These forms enforce mandatory fields, predefined data types, and controlled vocabularies. Real time validation is applied during data entry, allowing inconsistencies, such as missing fields or non-standardized terminology, to be identified and corrected.

**Figure 3:**
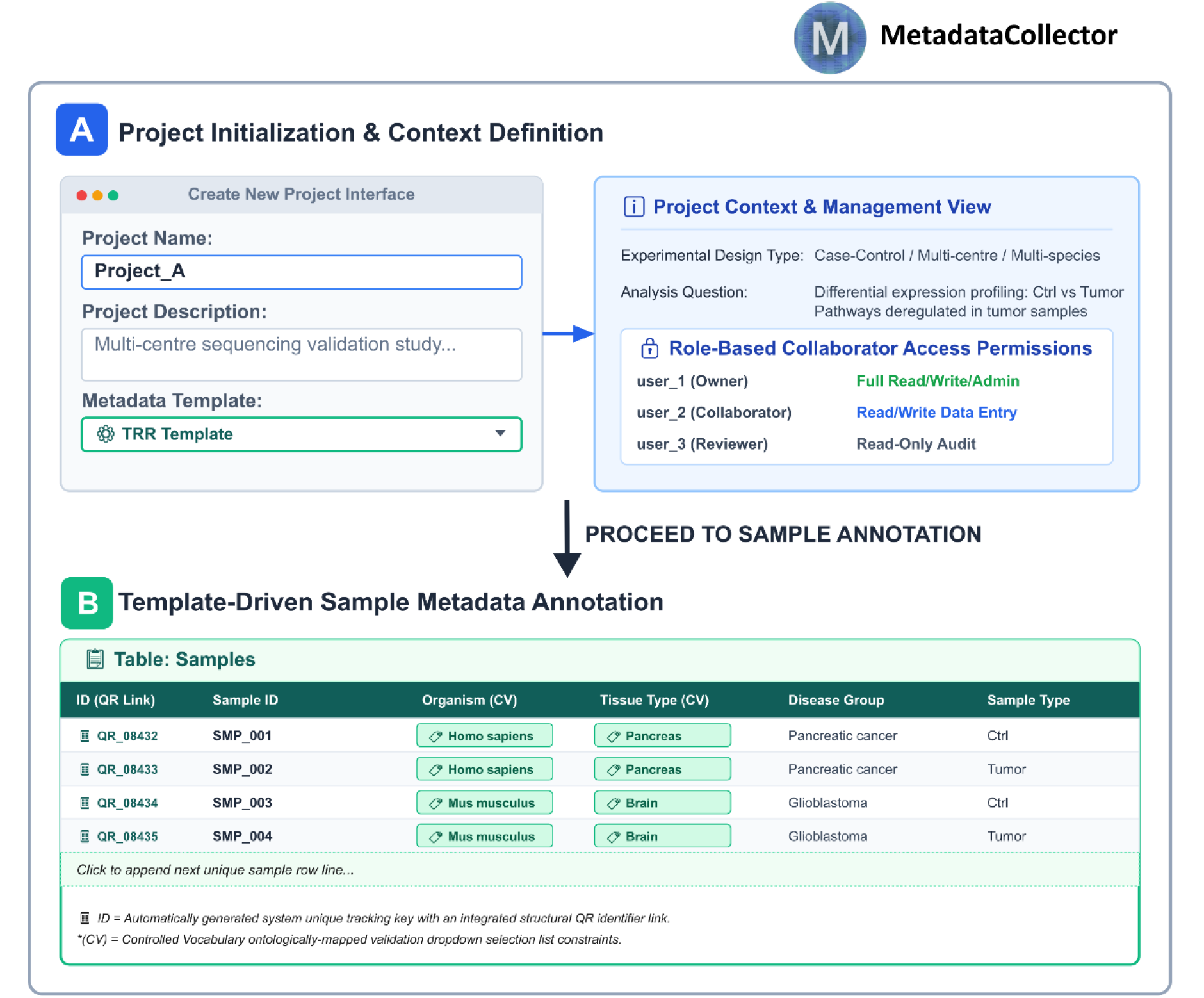
Project creation and sample annotation workflow in Metadata Collector. (A) Project initialization captures study-level information, metadata templates, and role-based collaborator permissions. (B) Biospecimen metadata are recorded in a structured annotation table containing unique sample identifiers and controlled vocabulary-supported fields. Together, these components facilitate standardized, validated, and FAIR-aligned metadata collection

Project administrators can modify templates during the course of the study using a dedicated configuration interface. Study-specific fields, such as treatment conditions or extraction protocols, can be added to the template, with updates propagated automatically to all project members without affecting existing records.

All metadata modifications are captured through an event-based versioning mechanism that records timestamps and contributor identities. This enables full provenance tracking and supports transparent review of project history (Figure 2). Sample-level descriptors such as organism, tissue type, disease group, and library strategy are populated using controlled vocabulary fields, while optional free-text fields allow additional experimental context to be documented when necessary.

Upon completion of the project, harmonized metadata can be exported in repository-oriented formats, including CSV and ISA-Tab, for downstream analysis and submission (Figure 3). The exported files contain complete mandatory fields, standardized terminology, and project-level descriptors aligned with repository requirements, facilitating efficient repository deposition.

## Discussion

Accurate and consistent metadata collection and curation remain a major bottleneck in multi-centre sequencing projects, where information is typically collected across heterogeneous laboratories using spreadsheets or ad hoc templates. Metadata Collector addressed this challenge by shifting quality control upstream, enabling contributors to create, validate, and harmonize records collaboratively rather than relying on late-stage curation. The platform’s combination of customizable templates, controlled vocabularies, real time validation, and event-based versioning provides a practical framework for improving the completeness and consistency of metadata at the point of entry.

Compared with existing solutions such as ISA-Tab validators, submission portals, or institution-specific LIMS, Metadata Collector occupies a complementary niche: it is not a submission tool or a laboratory tracking system, but an intermediate layer that facilitates harmonized metadata creation before downstream processing. This proactive approach reduces the iterative correction cycles commonly encountered in repository submissions and improves interoperability by enforcing structured fields aligned with community standards and repository requirements. The use case and adoption data from the TRR 305 consortium illustrate how early validation, shared templates, and vocabulary constraints can reduce terminology heterogeneity and streamline multi-group communication.

The underlying architecture and workflows are generic and transferable to other multi-omics research consortia, institutional sequencing cores, and collaborative clinical research networks that face similar challenges in distributed metadata creation and harmonization. This transferability is supported by the separation of frontend, backend, and data model components, which enables deployment with minimal customization, as well as by domain-agnostic design choices such as the ontology-ready vocabulary model, the flexible template system, and the event-sourcing mechanism. These characteristics allow Metadata Collector to be adapted to a broad range of research environments without requiring substantial changes to the underlying architecture. However, the platform is not intended to replace repository submission systems or full LIMS solutions. Instead, it complements these infrastructures by providing a collaborative metadata harmonization layer during project execution, helping to ensure that metadata are standardized, validated, and traceable before downstream analysis or repository submission.

At the same time, several limitations should be acknowledged. First, the assessment of platform impact was based primarily on adoption metrics, curator feedback, and practical use within the consortium rather than on formal usability studies or systematic benchmarking. Second, although real time validation and controlled vocabularies help reduce annotation inconsistencies, their effectiveness depends on continued template maintenance and vocabulary curation. Finally, while performance remained stable for projects comprising up to approximately 700 samples in our deployment environment, scalability to substantially larger datasets warrant further evaluation. Future work will focus on expanding quantitative assessment of platform performance and usability, as well as supporting increasingly large and diverse multi-omics projects.

## Conclusion

Metadata Collector provides a practical and scalable approach for improving the quality, consistency, and traceability of metadata in multi-centre NGS projects. By enabling contributors to work with shared templates, controlled vocabularies, and real time validation, the platform shifts metadata harmonization upstream, reducing the need for downstream curation and correction. Its event-based versioning model and secure, role-aware architecture support transparent collaboration and robust provenance tracking throughout the metadata lifecycle.

The successful deployment of Metadata Collector within the TRR305 consortium demonstrates its applicability in real-world multi-omics research environments. Moreover, its modular architecture, flexible template system, and ontology-ready design make it adaptable to a wide range of research settings beyond the consortium in which it was developed. Future extensions, including deeper ontology integration, expanded interoperability with external systems, and formal usability evaluation, will further strengthen its capabilities. As biomedical research increasingly relies on collaborative, multi-institutional data generation, tools that support standardized and FAIR-aligned metadata management will play an increasingly important role in ensuring data quality, reproducibility, and long-term reusability.

## Availability & Reproducibility

Software versions and deployment instructions are documented and versioned in the public GitHub repository. https://github.com/spang-lab/metadata-collector

## Ethics / Data Protection

Metadata managed in Metadata Collector are pseudonymized and do not include direct identifiers; therefore, no additional ethical approval for this manuscript was required.

## Acknowledgments

We thank all members of the TRR305 consortium for their valuable contributions and constructive input during the development and evaluation of Metadata Collector. This study was supported by the Deutsche Forschungsgemeinschaft (DFG, German Research Foundation) project TRR 305-Z01 to F.F and project 509149993, TRR 374-INF to F.F.

## Notes

https://github.com/spang-lab/metadata-collector

### Competing Interest Statement

The authors have declared no competing interest.

### Summary of Updates

This version of the manuscript has been revised to update the following: Author affiliations; funding.

